# Disrupting short-term memory in premotor cortex affects serial dependence in visuomotor integration

**DOI:** 10.1101/2021.02.18.431802

**Authors:** Raymundo Machado de Azevedo Neto, Andreas Bartels

**Affiliations:** Brain Institute, Hospital Israelita Albert Einstein, São Paulo 05652- 900, Brazil; Max Planck Institute for Biological Cybernetics, Tübingen, Germany; Centre for Integrative Neuroscience, Tübingen, Germany; Department of Psychology, University of Tübingen, Tübingen, Germany; Bernstein Center for Computational Neuroscience, Tübingen, Germany

**Author notes:** Corresponding author: Raymundo Machado de Azevedo Neto.

## Abstract

Human behavior is biased by past experience. For example, when intercepting a moving target, the speed of previous targets will bias responses in future trials. Neural mechanisms underlying this so-called serial dependence are still under debate. Here, we tested the hypothesis that the previous trial leaves a neural trace in brain regions associated with encoding task-relevant information in visual and/or motor regions. We reasoned that injecting noise by means of transcranial magnetic stimulation (TMS) over premotor and visual areas would degrade such memory traces and hence reduce serial dependence. To test this hypothesis, we applied bursts of TMS pulses to right visual motion processing region hV5/MT+ and to left dorsal premotor cortex during inter-trial intervals of a coincident timing task performed by twenty healthy human participants (15 female). Without TMS, participants presented a bias towards the speed of the previous trial when intercepting moving targets. TMS over dorsal premotor cortex decreased serial dependence in comparison to the control Vertex stimulation, whereas TMS applied over hV5/MT+ did not. In addition, TMS seems to have specifically affected the memory trace that leads to serial dependence, as we found no evidence that participants’ behavior worsened after applying TMS. These results provide causal evidence that an implicit short-term memory mechanism in premotor cortex keeps information from one trial to the next, and that this information is blended with current trial information so that it biases behavior in a visuomotor integration task with moving objects.

**Significance Statement:** Human perception and action are biased by the recent past. The origin of such serial bias is still not fully understood, but a few components seem to be fundamental for its emergence: the brain needs to keep previous trial information in short-term memory and blend it with incoming information. Here, we present evidence that a premotor area has a potential role in storing previous trial information in short-term memory in a visuomotor task, and that this information is responsible for biasing ongoing behavior. These results corroborate the perspective that areas associated with processing information of a stimulus or task also participate in maintaining that information in short-term memory even when this information is no longer relevant for current behavior.

## Introduction

When executing motor tasks that depend on visual information, the central nervous system combines sensory with prior information (e.g., Kording and Wolpert 2004). In particular, information from the immediately preceding trial can affect behavior (Kowler et al. 1984; de Lussanet et al. 2001; Gray 2002; Jarrett and Barnes 2002; Heinen et al. 2005; Makin et al. 2008; Kwon and Knill 2013). For instance, Kwon and Knill (2013) have shown that temporal error in an interceptive action is biased by the speed of the previous trial. Specifically, participants anticipate responses on trials preceded by faster target speeds, and delay responses on trials preceded by slower target speeds. Note that this effect is opposite to that expected by simple adaptation in visual or motor regions (c.f. Thompson and Burr, 2009). Also, this effect is not exclusive to visuomotor tasks, being also reported in visual perception, where it was dubbed serial dependence (Fischer and Whitney, 2014). The mechanism supporting the serial dependence effect is still under debate.

One possible explanation for serial dependence in visuomotor integration tasks is that information from the previous trial blends with incoming sensory and/or motor planning information (de Lussanet et al. 2001; Makin et al. 2008; Kwon and Knill 2013). For instance, interception response bias might emerge from estimating target speed using a weighted average between previous and current target speeds. For this blending to occur, the previous trial speed should be stored during the inter-trial interval. Therefore, serial dependence would rely on a short-term memory mechanism.

If serial dependence in visuomotor integration tasks relies on a short-term memory mechanism, which brain regions store information from the previously experienced trial? One prominent proposal is that those cortical regions involved in processing and encoding the specific sensory and motor information of a specific task also participate in its short-term maintenance (for reviews, see Postle 2015; Serences 2016; Christophel et al. 2017). Performing a visual-motor interception task has been shown to engage many regions along the visual and motor cortices (Indovina et al. 2005; Miller et al. 2008; Maffei et al. 2010; de Azevedo Neto and Amaro Jr, 2018). Hence, storing previous trial information in-between trials might involve both visual as well as motor planning regions. To support this idea, imaging evidence has shown visual and motor regions to store short-term memory, although in explicit short-term memory tasks (Christophel et al. 2012; Jerde et al. 2012; Riggall and Postle, 2012; Christophel and Haynes 2014).

Having such a distributed short-term memory storage framework in mind (c.f. Christophel et al. 2017), we hypothesized that serial dependence in a visuomotor task would rely on memory traces implicitly stored in regions involved with the task along the visuomotor processing hierarchy.

Specifically, we tested for causal contributions of two sites of the processing stream: serial dependence might be caused by early visual motion representations in motion region V5/MT+, or upstream motor planning information in premotor cortex. We used a simple interceptive task (i.e. coincident timing task) to study this phenomenon. For this task, we expected that either premotor or a visual motion sensitive region would encode information from the previous trial during the gap between trials, since these areas have been associated with performing interceptive tasks (Indovina et al. 2005; Miller et al. 2008; Maffei et al. 2010; de Azevedo Neto and Amaro Jr, 2018). We acknowledge, though, that many other regions related to motor planning and execution (c.f. Connolly et al. 2007) as well as other areas sensitive to visual motion (c.f. Orban et al. 2003) could also be responsible for keeping information in short-term memory from one trial to the next, and that our experimental design would not allow testing for all possible regions along the visuomotor processing hierarchy. We assumed that injecting noise by means of transcranial magnetic stimulation (TMS) in-between trials would degrade such short-term memory traces by removing the specific tuning in synaptic weights left by the specific stimulus that was just experienced (Barbosa et al. 2020; Bliss and D’Esposito 2017; Kilpatrick 2017; van de Ven and Sack 2013; Zokaei et al. 2014) and hence reduce serial dependence.

## Materials and Methods

### Participants

Twenty healthy young adults (15 female, 5 male; mean age 25.5 years, standard deviation 3.76 years) participated in the reported experiment. All participants were right-handed, had normal or corrected to normal vision, and had no contraindications for TMS (Rossi et al. 2009) or fMRI. Participants provided written informed consent prior to the study. The study was approved by the Local Ethics Committee of the University Clinic Tübingen.

Participants received monetary compensation for participation.

### Overall procedure

In the first session, participants were screened for their suitability for TMS (Rossi et al. 2009) and fMRI experiments. Also on the first session, participants performed the coincident timing task without TMS. In the second session, they then underwent an fMRI localizer experiment that served to identify their individual visual motion regions V5/MT+ as well as the dorsal Premotor Cortex (PMd), using separate sets of stimuli and task conditions described below in detail. In the third and last session, following fMRI localizers, participants took part in the TMS experiment (Figure 1A). Below, we first describe stimuli, task and analyses of the TMS experiment, followed by those of the fMRI localizers.

**Figure 1.**
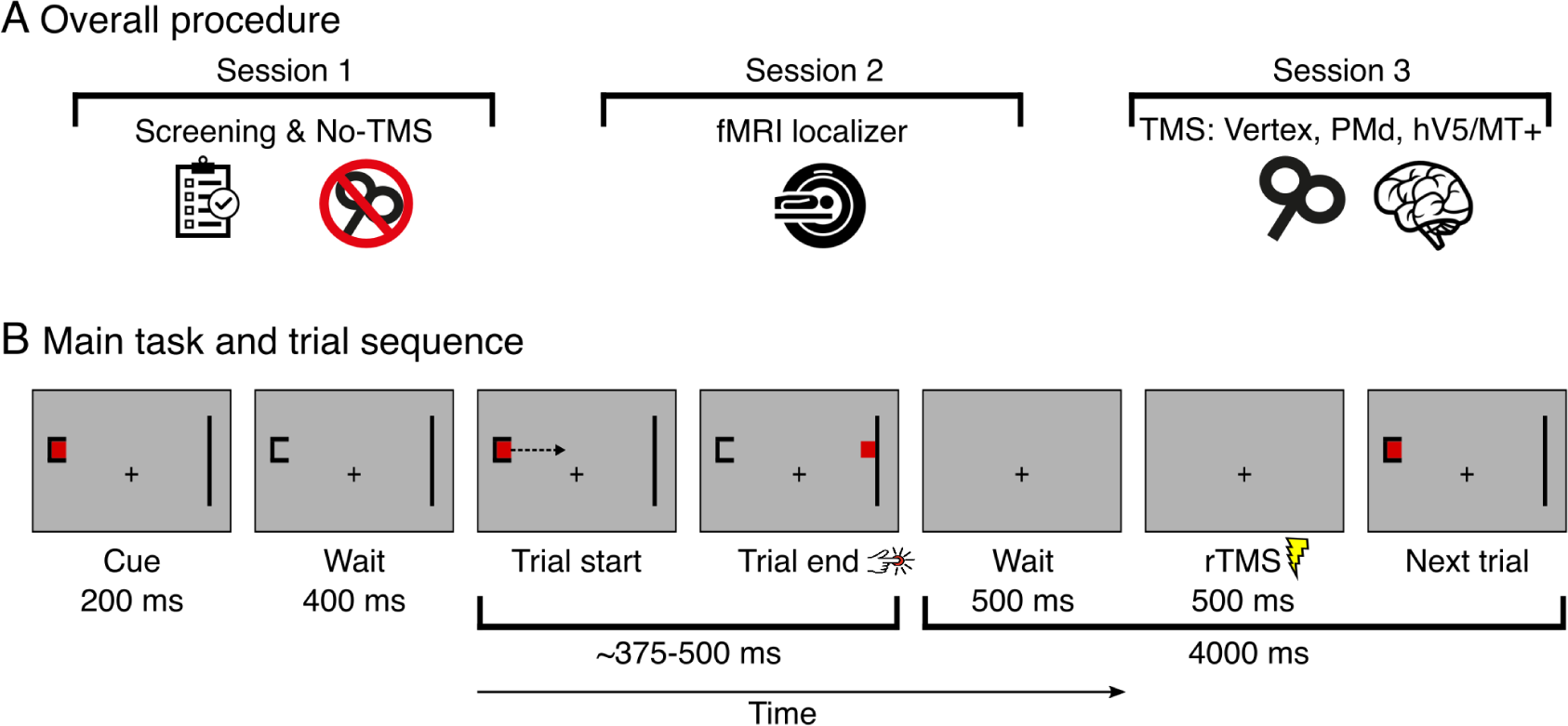
Overall procedure and TMS trial schematics. (A) In three separate sessions, participants were screened for TMS and fMRI suitability (session 1), performed an fMRI localizer experiment to individually determine TMS target sites (session 2), and performed the main coincident timing task while we applied TMS at hV5/MT+, PMd, or Vertex (session 3). (B) On each trial of the main task on the first and on the TMS sessions, participants were presented a target moving rightwards and were required to press a button at the same time as the target hit the interception zone. We applied TMS bursts 500 ms into the pause between trials.

### TMS: Stimuli and task

Participants performed a coincident timing task while seated in a comfortable chair and with their head positioned on a chin rest at 57 cm from an LCD screen (14.1 inches, 1600 x 1200 pixels, 60 Hz). Stimuli and task were generated and controlled using MATLAB (http://mathworks.com/) and Psychtoolbox (http://psychtoolbox.org/). Throughout the experiment, participants were instructed to fixate their gaze at a cross on the center of the screen. The fixation cross was present on the screen for the entire experiment right below the target’s motion path (see below). Eye tracking was performed throughout the experiment at 500 Hz (see separate section below for details). On each trial, participants saw a red square target (0.5° of visual angle) that moved from left to right of the screen at a constant speed (20, 22, 24, 26, 28°/s). The motion trajectory always spanned 10° of visual angle, resulting in motion times of 0.357 to 0.5 seconds. The beginning of each trial was cued by presenting the start zone (squared black letter “C”) on the left side of the screen and the interception zone (vertical black bar) on the right side of the screen. At the same time as the start and interception zones were presented, the target was flashed for 200 ms to warn participants that the trial was about to begin. After 400 ms, the target was back on the screen, but now moving from left to right (Figure 1B). Participants were instructed to press a button at the exact moment the target would hit the bar at the end of its trajectory (button box sampling frequency 1000 Hz). After 500 ms of target arrival at the interception zone, the start and interception zones vanished from the screen. Then, a pause lasting 4 s occurred until the onset of the next trial. Note that fixation was required also during the pause.

### TMS: Experimental design and procedures

To evaluate the serial dependence effect, we need to prevent that participants learn a particular sequence of targets, guarantee the same transition probabilities across target speeds and guarantee the same number of trials for each target speed. To avoid these confounders, all combinations of previous and current trial speeds were presented the same number of times, with target speeds having the same probability of being preceded by every of the 5 target speeds presented in the experiment (Brooks 2012).

Participants familiarized with the task by performing 20 trials before the main experiment. Each speed was randomly presented four times during practice. During the main experiment, participants performed the task in four conditions (described below). All conditions were performed in 5 consecutive blocks, each with 51 trials. Target speed counterbalancing was performed independently for each block. The first trial of each block was discarded from further analyses. Each combination of previous and current trial speeds (25 possibilities) was repeated twice per block, amounting to 10 trials per pair of speeds. Between each block, participants were allowed to rest for 1 min, as informed by a clock on the center of the screen. During this rest interval, participants were asked to keep their heads still on the chin rest. Participants took ∼ 4 min to complete each block, and ∼ 20 min to complete each condition (excluding resting intervals).

Participants performed the task under 4 conditions. In one of them (first session) the task was performed without any stimulation (No-TMS condition). On the other 3 conditions (third session), TMS pulses were applied during the gap between trials on right hV5/MT+, left PMd, or Vertex (Figure 2C). On TMS stimulation conditions, the pulses started 500 ms after the end of a trial, leaving enough time for dissipating the rTMS effect and not affecting visual and motor processing on the next trial (Walsh and Pascual-Leone 2003; Campana et al. 2007; Bosco et al. 2008). The order of TMS stimulation conditions was counterbalanced across participants.

**Figure 2.**
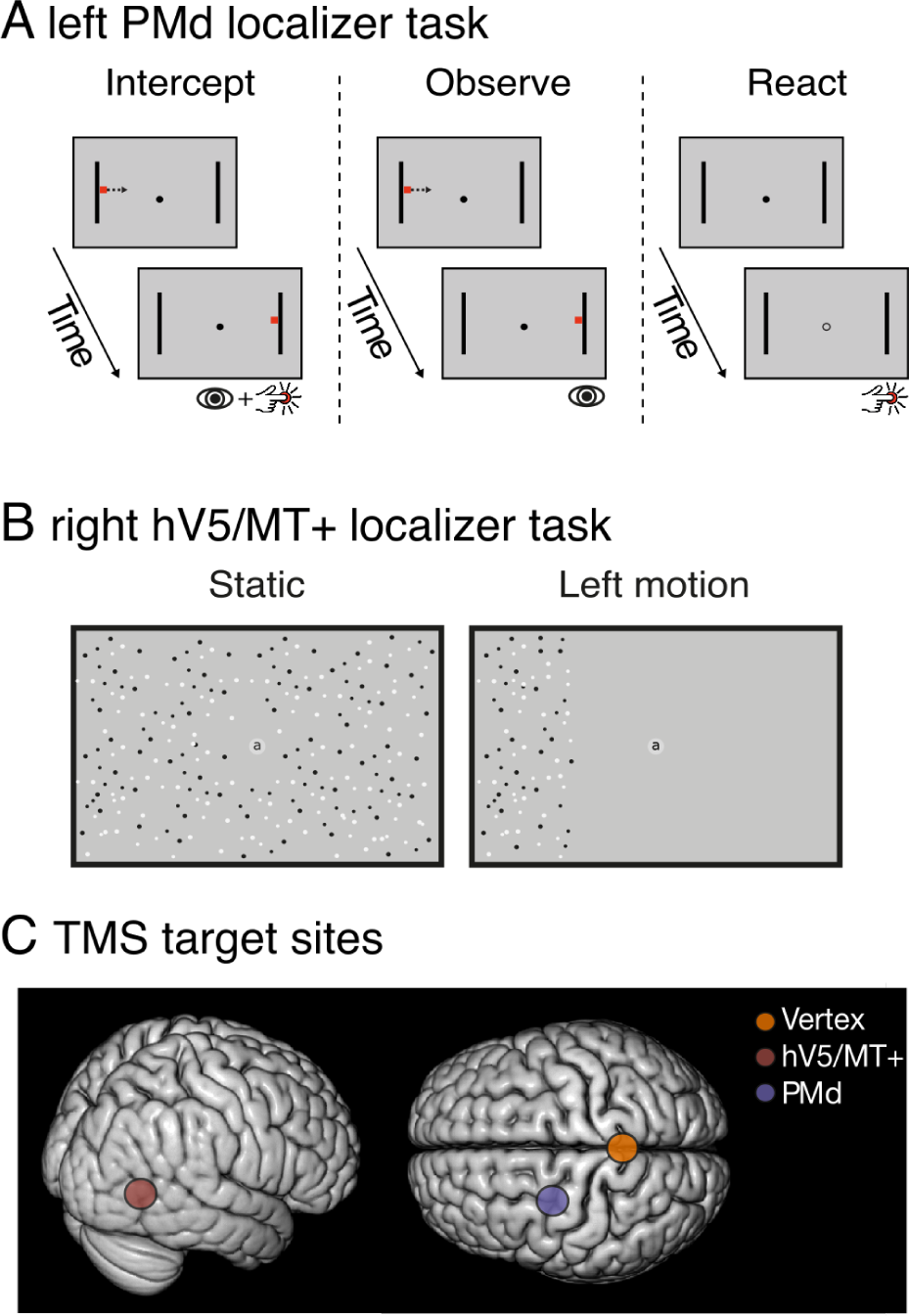
Stimulus conditions of the functional localizer tasks and illustration of TMS stimulation sites. Left PMd and right hV5/MT+ were individually defined using fMRI in a separate localizer experiment preceding the TMS experiment. (A) Left PMd was defined as the region showing increased BOLD signal when intercepting moving targets (left) contrasted against only viewing moving targets (middle) and performing a reaction time task (right). (B) Right hV5/MT+ was defined as the region showing increased BOLD signal when participants viewed moving dots (left) at the left visual hemifield contrasted with static moving dots (right). (C) Illustration of TMS stimulation sites at right hV5/MT+, left PMd, and Vertex. Vertex was defined individually on the anatomical images.

Participants had a 15 min rest period between TMS stimulation conditions.

### TMS: Behavioral data analysis

We evaluated participants’ behavior by measuring their temporal error using custom made scripts in MATLAB and further statistical analyses and figures were performed in R (version 2.15.2) and JASP (0.9.2). Temporal error was defined as the difference between the moment the target hits the interception zone and the moment participants press the button. Positive values indicate participants were late and negative values indicate they anticipated their responses.

For each stimulation condition and for each participant, we discarded data from trials in which the absolute temporal error was higher than three times the standard deviation for that specific condition and participant (for similar procedures, see Makin et al. 2008; Kwon and Knill 2013). In addition, we excluded trials in which participants pressed the button 500 ms before or after target arrival at the interception zone. Using this exclusion criterion, 4.94%, 3.8%, 2.48%, and 3.32% of trials were excluded from No-TMS, Vertex, left PMd and right hV5/MT+ conditions, respectively. All data from one participant was excluded from further analyses because the participant failed to comply with the task instruction by pressing the button too late on average, what would be expected in a reaction time task. One participant did not complete left PMd condition because of a headache and we excluded data from this participant on this condition. Completely removing this participant from the sample did not change the results from the analyses.

Data from one participant at No-TMS condition was excluded because the button box failed to record more than half of this participant’s responses.

We modeled participants’ temporal error by means of a multiple linear regression model:

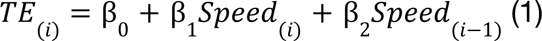

where TE is the temporal error on trial *i*, β_0_ is a constant bias term, and the independent variables *Speed_(i)_* and *Speed_(i-1)_* are speeds for current and previous trials, respectively. This equation was used to estimate parameters β_1_ and β_2_ for each participant in each of the stimulation conditions separately.

In our experiment, serial dependence would be detected by observing negative values for β_2_ in our linear regression, which would indicate that the higher the previous target speed, the greater anticipation bias in participants’ responses. To test this, we performed a one-tailed one-sample Wilcoxon signed rank test for the estimated values of β_2_ from all participants at the No-TMS condition. More importantly, to check whether applying TMS over hV5/MT+ or PMd decreases serial dependence, we performed one-tailed Wilcoxon signed rank tests for β_2_ values in these two conditions against Vertex control condition. A decrease in serial dependence would be detected by an increase in β_2_ values toward zero. Our hypothesis is that applying TMS over regions that putatively keep a memory trace from the previous trial will decrease serial dependence. We assume that these regions might keep information through a specific tuning in synaptic weights that were strengthened after experiencing the previous trial (Barbosa et al. 2020; Bliss and D’Esposito 2017; Kilpatrick 2017). After applying TMS over such regions of the cortex, the electromagnetic pulse should instantaneously make many neurons fire at once, thus decreasing or removing the specific tuning left by the previous trial stimulus or action. This specific and directional hypothesis will correspondingly be tested using a one-tailed test. To verify that serial dependence was still present after applying TMS over right hV5/MT+, left PMd and Vertex, we performed one-tailed one sample Wilcoxon signed rank tests on β_2_ values.

If applying TMS over stimulation sites during the gap between trials had unintended effects such as distracting participants or affecting sensorimotor processing of the next stimulus and response, we should observe differences in the slope of current trial speed for the multiple linear regression analysis (β_1_ values) or on the overall temporal error across conditions, as well as increased variability in participant’s responses in stimulation conditions. To check for these effects, we submitted β_1_ values, overall temporal error and variable error (standard deviation of temporal error) values aggregated across all speeds for each condition to Friedman test for repeated measurements. Significance level (α) was set at 5% for all statistical analyses of behavioral data and applied Bonferroni correction for multiple comparisons within each family of tests addressing a specific research question using. We defined the following families of tests where correction for multiple comparisons were applied: 1) presence of serial dependence effect on each condition (one-sample tests on β_2_); 2) effect of TMS over the cortex on the serial dependence effect (paired tests on β_2_); 3) effect of TMS over the cortex on the current trial bias (paired tests on β_1_).

### rTMS protocol

Biphasic rTMS (10 Hz, 6 pulses = 500 ms) were applied 500 ms after the beginning of the interval between trials using an eight shaped TMS coil, with 97 cm external diameter (MC-B70) connected to a MagPro X100 stimulator (MagVenture, Farum, Denmark). rTMS pulses were automatically triggered using the same scripts that controlled stimuli presentation.

Stimulation parameters used were in accordance with safety guidelines (Rossi et al. 2009).

After task familiarization, neuronavigation calibration was performed to help position the coil over participant’s head (TMS-Navigator, Localite, Sankt Augustin, Germany). Calibration consisted of registering physical coordinates of participants’ heads acquired with infrared cameras into participants’ T1-weighted images.

TMS intensity was estimated for each participant based on their resting motor threshold after localizing the hot spot of the right hand region in the left primary motor cortex. To assist in finding the hot spot, we identified the left hemisphere hand region on the T1-weighted images before the experimental session and positioned the TMS coil over it. Hot spot was identified as the region that clearly elicited finger movements with single pulses around the predetermined hand region. Initial stimulation intensity for all participants was set to 35% of the maximum output of the stimulator.

When necessary, stimulation intensity was increased in steps of 2% of the maximum output of the stimulator. Resting motor threshold was defined as the lowest single pulse stimulation intensity that elicited 5 out of 10 movements in a sequence. The TMS coil was positioned tangentially to the scalp of participants with a 45° angle with respect to horizontal and with handle pointing back. During the experiment, the intensity for each participant was set as 100% of resting motor threshold. Mean intensity used in the experiment was 39.95% (± 3.72 standard deviation) of maximum output of the stimulator.

### TMS: Eye-tracker data acquisition and analysis

Eye movements were recorded using an Eyelink 1000 Plus Desktop system (SR Research Ltd., Mississauga, Canada) at 1000 Hz. The eye tracker was calibrated prior to each stimulation condition by having participants fixate 9 targets at known eccentricities. Eye movements were analyzed using custom made software in MATLAB. Eye blinks were identified and data before (100 ms) and after (100 ms) blinks were removed and linearly interpolated. Time series were filtered using a recursive 4th order low-pass Butterworth filter with cut-off frequency of 30 Hz, and linear trend was removed. Trials with values that are out of the screen boundaries were excluded from further analyses. Drifts in eye position recording that occur throughout the experiment were corrected offine by first estimating the mode of the eye position cluster during the inter-trial interval preceding each trial, then calculating the distance between this mode and the actual fixation cross position, and finally subtracting this distance from the time series during the trial period (Zhang and Hornof 2011). On each trial, we identified the number of saccades by detecting eye movements with velocity higher than 30°/s and displacements greater than 2° visual angle. The velocity time series was estimated using a two-point difference algorithm. To control that participants’ eye movements did not differ across conditions, we estimated the number of saccades and percentage of time that gaze was within 2° around the fixation cross. These data were submitted to a one-way repeated measures ANOVA.

### fMRI localizer and definition of TMS stimulation sites

We defined TMS stimulation sites for each participant in the second session using fMRI localizer tasks combined with anatomical images. Table 1 presents the mean (± standard deviation) center of mass for stimulation sites across participants. During the fMRI experiments, participants saw stimuli through a mirror mounted on top of the head coil, which reflected the projected image (Projector HP, 1280 x 1024 pixels, 60 Hz) on an acrylic screen at the end of the scanner bore. Visual display subtended 24° x 18° of the visual field. Participants’ responses on both tasks were acquired using an MR compatible button box, with sampling frequency of 1000 Hz. All stimuli, behavioral measures and MR synchronization were performed using MATLAB and Psychtoolbox.

**Table 1.** MNI coordinates (mean ± standard deviation) for the three stimulation sites (right hV5/MT+, left PMd, and Vertex).

| Stimulated region | X | Y | Z |
| --- | --- | --- | --- |
| Right hV5/MT+ | 45.47 ± 6.24 | -64.62 ± 4.42 | 8.42 ± 5.53 |
| Left PMd | -26.00 ± 6.92 | -4.48 ± 7.22 | 54.73 ± 5.54 |
| Vertex | -0.21 ± 1.61 | -37.47 ± 6.14 | 92.42 ± 3.02 |

### fMRI: Left PMd localizer task

To individually identify left PMd, participants performed an fMRI experiment that has previously identified this region in an interceptive task similar to that used in the main experiment (de Azevedo Neto & Amaro Jr, 2018). We chose to stimulate left PMd because participants performed the main experiment with their right hand.

Before entering the scanner, participants practiced and familiarized with the task for approximately 1 min. Participants performed an event-related experiment with three conditions: Intercept, Observe and React (Figure 2A). Before each condition, participants were warned about the condition on the next trial by presenting a word corresponding to the condition for 500 ms. After the instruction disappeared, two vertical bars (3.5° x 0.33° visual angle, height x width, black) and a fixation dot (0.33° visual angle diameter) were presented. In the Intercept condition, as soon as the trial started, a red square target (0.5° of visual angle) appeared on the screen beside one of the vertical bars and started moving towards the opposite vertical bar. Participants were instructed to press a button with their right thumb at the exact same time the target reached the opposite vertical bar (Figure 2A, left panel). The target took between 370 ms and 833.33 ms (370, 416.66, 476.19, 555.55, 666.67, 833.33 ms) to travel across the screen. In the Observe condition, participants were required to pay attention to target motion displacement, without any motor response (Figure 2A, middle panel). Target traveling time was the same in the Observe and Intercept conditions.

In the React condition, participants were required to press a button as soon as they perceived the fixation dot had transformed into a hollow black ring (Figure 2A, right panel). The transformation of the fixation dot could occur between 370 ms and 833.33 ms (370, 416.66, 476.19, 555.55, 666.67, 833.33 ms) after the instruction disappeared from the screen. At the end of each trial, the vertical bars were removed from the screen. Inter-trial interval was randomly chosen from a uniform distribution between 3 and 5.5 s

Conditions were counterbalanced with the restriction that each condition had the same probability of being preceded by all conditions (Brooks 2012). Participants performed 42 trials per condition, with 7 trials for each of the possible timing parameters presented above. Target motion direction in Intercept and Observe conditions was randomly chosen.

Participants did not receive any feedback about their performance throughout the experiment. This experiment run took approximately 11 min to be completed.

### fMRI: hV5/MT+ localizer task

Functional localization of right hV5/MT+ was performed by presenting moving stimuli to the left visual hemifield of participants (motion condition) and comparing this with static images of the same stimuli (static condition), a common task used in the literature for the purpose of defining visual motion areas (Zeki et al. 1991; Huk et al. 2002; Fischer et al. 2012)(Figure 2B). We chose to stimulate right hV5/MT+ because we had no a priori hypothesis about differences in short-term memory function across cerebral hemispheres. Since targets were presented at 5° of visual angle to both sides of the fixation cross, this allowed area hV5/MT+ of both hemispheres to receive visual stimulation (Dukelow et al. 2001; Huk et al. 2002).

In the static condition, participants saw 1540 black and white dots randomly spread across the screen (100% contrast, diameter between 0.1° to 1.1° in visual angle) in a gray background (dot density 0.75 dots/°^2^, 90 cd/m^2^). In the motion condition, participants saw on average 1540 black and white dots spread across 3/5 of the left hemifield from the left border of the screen. All dots moved coherently and simultaneously while keeping their relative distances fixed. Each condition was presented 7 times in blocks of 12 s. In the same run, participants performed another 5 conditions, each one with 12 s, but these were not used for the purposes of the present study and are not reported here. All conditions were presented in a pseudo-random order, with the restriction that each condition had the same probability of being preceded by all conditions (Brooks 2012). The experimental run lasted approximately 10 min.

In both conditions, participants performed a fixation task to guarantee they would keep their gaze on a gray disc and balance attentional load across conditions. In this task, a sequence of letters was presented and participants were instructed to press a button when detecting letter repetitions. Every three to eight presentations of one of these characters were repeated. A total of 26 characters were presented randomly (1.6° height in visual angle, black) projected inside a gray disc (2° visual angle in diameter, 72 cm/m^2^) at the center of the screen.

### fMRI acquisition

We acquired functional T2*-weighted parallel gradient-echo multiplexed echoplanar images (EPI) on a Siemens MAGNETON Prisma 3 T scanner using a 64-channel phased-array coil. The pulse sequence included multiband radio frequency excitation with factor 2 and were acquired with the following parameters: TR = 1200 ms, TE = 30 ms, flig angle = 68°, FOV = 192 x 192 mm, slice thickness = 3 mm, acquisition matrix = 64 x 64, 36 slices without gaps, voxel size = 3 x 3 x 3 mm. We recorded 580 volumes for the PMd localizer task, and 515 volumes for the hV5/MT+ localizer task. The first 8 volumes on each run were acquired in the absence of any task to allow for signal stabilization and excluded from further analyses. In addition, a high-resolution anatomical scan was also obtained for each participant with a T1-weighted MP-RAGE sequence (ADNI, 192 slices, 1 x 1 x 1 mm, TR = 2000 ms, TE = 3.06 ms, FOV = 232 x 256 mm).

### fMRI: Preprocessing

fMRI data processing was performed using FEAT (FMRI Expert Analysis Tool) Version 6.00, part of FSL (FMRIB’s Software Library, www.fmrib.ox.ac.uk/fsl). All preprocessing procedures were the same for both localizer runs, except otherwise stated. Functional images were registered in the T1-weighted anatomical image using FLIRT (Jenkinson and Smith 2001; Jenkinson et al. 2002). The following sequence of preprocessing steps was performed: motion correction using MCFLIRT (Jenkinson et al. 2002); slice-timing correction using Fourier-space time-series phase-shifting; non-brain removal using BET (Smith 2002); spatial smoothing using a Gaussian kernel of FWHM 5 mm; grand-mean intensity normalization of the entire 4D dataset by a single multiplicative factor; high-pass temporal filtering (Gaussian-weighted least-squares straight line fitting, with sigma = 64 s for hV5/MT+ localizer task and sigma = 50 s for PMd localizer task). Time series statistical analysis was carried out using FILM with local autocorrelation correction (Woolrich et al. 2001).

### fMRI statistical analyses

Statistical analysis was implemented using a massive univariate approach using a General Linear Model on each voxel for both localizer tasks.

### fMRI statistical analysis: left PMd localizer task

Activity was modeled on each trial as a single event. The beginning of each trial was determined as the moment the target appeared on the screen and started moving. The end of each trial was determined as the moment the target arrived at the opposite vertical bar in its trajectory or the time it took for the fixation dot to transform into a ring. Each condition — Intercept, Observe, and React — was modeled with one regressor and they were independently convolved with a double-gamma function representing the hemodynamic response function. Additional regressors modeled the estimated motion parameters for nuisance regression. Left PMd was individually identified as voxels at the intersection of the right superior frontal sulcus and right precentral sulcus (Bestmann et al. 2005; Davare et al. 2006, 2015; Oshio et al. 2010; Moisa et al. 2012; Stadler et al. 2012; Wymbs and Grafton 2013; Parmigiani et al. 2015; Fujiyama et al. 2016) responding more to the Intercept condition than to both Observe and React conditions. This analysis was performed by means of a conjunction analysis (Nichols et al. 2005) after performing one-sample t tests for the [Intercept > Observe] and [Intercept > React] contrasts. The statistical map of the conjunction analysis was not corrected for multiple comparisons and thresholded flexibly (Z values at least greater than 2.3) in order to increase the chance of identifying left PMd for all participants and isolating it. In cases where the conjunction analysis did not reveal any voxels above the minimum uncorrected threshold, we used the one sample t-test contrast [Intercept > Rest], but still using the anatomical criteria described above (this happened for 12 out of 20 participants). Mean (± standard deviation) center of mass for left PMd is shown in Table 1.

### fMRI statistical analysis: hV5/MT+ localizer task

Activity in motion and static conditions was modeled as blocks of activity. One regressor for each condition was modeled as a boxcar of 12 s. Each regressor was convolved with a double-gamma function representing the hemodynamic response function. The remaining 5 conditions not reported here were modeled in the same way. Additional regressors modeled the estimated motion parameters for nuisance regression. Right hV5/MT+ was individually determined as voxels in the ascending limb of the inferior temporal sulcus (Dumoulin et al. 2000) responding more to motion than static stimuli in a one-sample t test. Results of this contrast were not corrected for multiple comparisons and thresholded flexibly (Z values at least greater than 2.3) in order to increase the chance of identifying right hV5/MT+ for all participants and isolating it (for similar approach, see Summerfield et al. 2008; Kovács et al. 2013). Because of technical problems during the localizer scan, we used only the anatomical criteria to determine right hV5/MT+ for one of the participants. Mean (± standard deviation) center of mass for right hV5/MT+ is shown in Table 1.

### Vertex identification

Vertex was defined on each participant’s T1-weighted anatomical images as the highest point on the scalp above the postcentral gyrus between both hemispheres. We decided to define Vertex at a different position compared to previous studies to avoid that TMS pulses would affect activity in the Supplementary Motor Area. Mean (± standard deviation) center of mass for Vertex is shown in Table 1.

## Results

If participants’ behavior is influenced by previous trial information, the temporal error on the current trial should be biased as a function of the target speed of the previous trial. We tested for such biases in the No-TMS and Vertex conditions. Figure 3 shows that for a given current trial speed, participants tended to give an earlier response when the previous trial speed was faster. Equally, participants tended to delay their responses when the previous trial speed was slower. In addition, the behavioral bias scaled linearly with the speed difference to the preceding trial: the greater the difference between current and previous trial speed was, the greater the bias turned out, as can be seen qualitatively by the color gradient across previous trial speeds (columns) for each current trial speed (rows) in Figure 3A, and in the scatterplots between previous trial speed and temporal error shown in Figure 3B. This indicates that participants anticipated or delayed their movements depending on the speed of the target on the previous trial and that this bias scales linearly at this range of target speeds.

**Figure 3.**
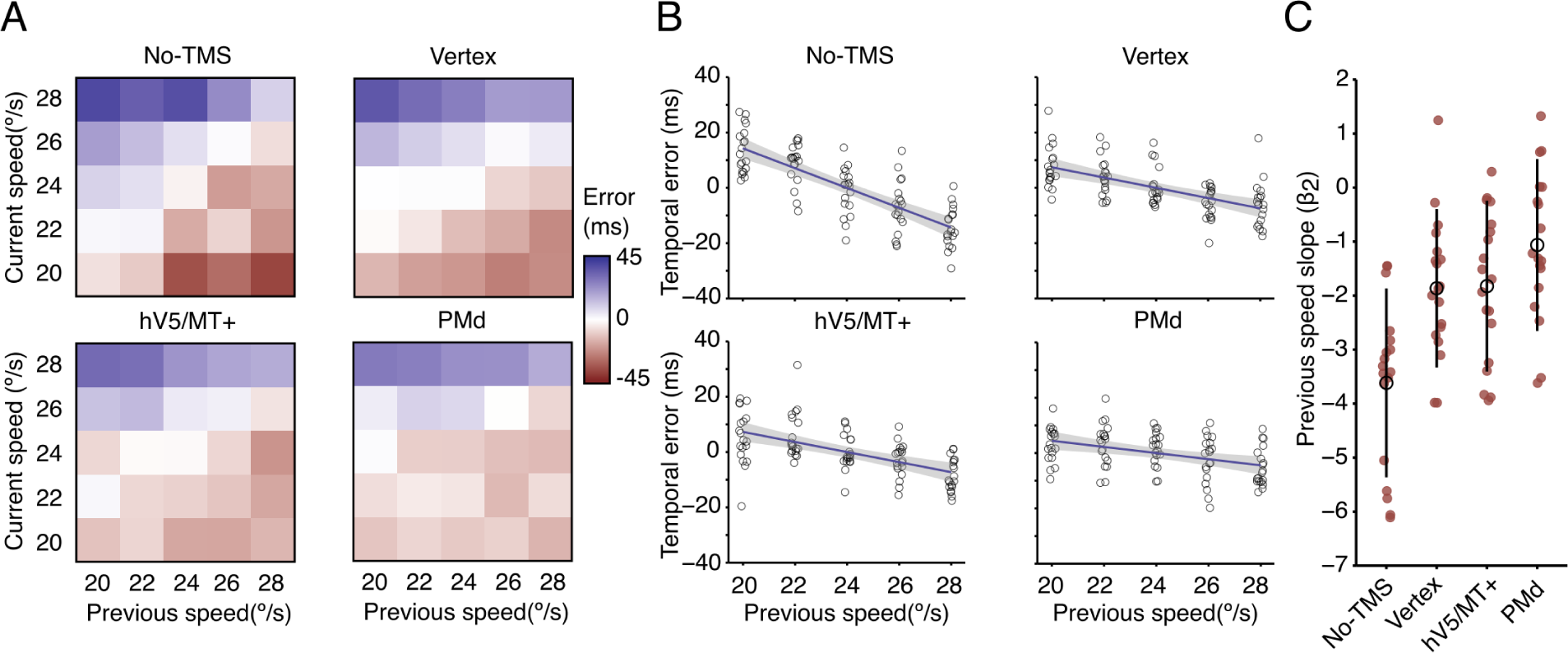
Behavioral results. (A) Temporal error as a function of previous and current trial speeds for No-TMS, Vertex, right V5/MT+, and left PMd conditions. Blue indicates positive temporal errors and red indicates negative temporal errors. (B) Linear regression for temporal error as a function of previous trial speed for No-TMS, Vertex, right V5/MT+, and left PMd conditions. Open circles represent the average temporal error for each participant at each previous target speed, also averaged over all current target speeds. Solid lines represent the best fitting line over all participants’ temporal errors, and shaded area represents the 95% confidence interval of the regression line. (C) Previous trial speed slopes of the multiple linear regressions for each participant at the No-TMS, Vertex, right V5/MT+, and left PMd conditions. Red circles represent slopes for multiple linear regressions of each participant, black open circles represent the median slope values across participants within each condition, and error bars represent the interquartile range.

To quantitatively assess this bias caused by the previous trial in the absence of TMS, we modeled the temporal error by means of a multiple linear regression taking into account the speed on current and previous trials (equation 1)(R^2^ values = median (25^th^ quantile, 75^th^ quantile); No-TMS = 0.102 (0.060, 0.193); Vertex = 0.061 (0.030, 0.178); PMd = 0.031 (0.006, 0.113); hV5/MT+ = 0.039 (0.022, 0.140)). The results show that the regressor for the previous trial is significantly different from 0 in the No-TMS condition [one-tailed one sample Wilcoxon signed rank test; V = 0; p_uncorrected_ < 0.001, p_bonferroni_ < 0.001, β = −3.62, Cohen’s *d* = 2.39] (Figure 3C). The negative slope indicates that the faster the speed of the previous trial, the more anticipated were the participants’ responses, as evidenced by more negative temporal errors on the current trial. These results are evidence for serial dependence in our interceptive action task.

If serial dependence relies on a short-term memory mechanism, which brain regions keep this information from one trial to the next? Based on prior literature, we hypothesized the same cortical regions involved with perceptual and motor processing of our interceptive action task would also keep the memory trace from previous trial, and causally contribute to serial dependence. We hence injected noise into these regions during the within the window separating our trials by applying TMS to either visual or motor regions. We expected that applying TMS over right hV5/MT+ or over left PMd would decrease the serial bias as a function of the previous trial in comparison to that observed in No-TMS and Vertex conditions. First, we verified that the previous trial speed slope was different across conditions [Friedman test; χ (3) = 19.8, p < 0.001, Kendall’s W = 0.388]. To quantify region-specific TMS effects, we then assessed whether regression slopes are bigger (less negative) for right hV5/MT+ and left PMd conditions than when applying TMS over Vertex. We found that slopes differed significantly from Vertex for left PMd [one-tailed Wilcoxon signed rank test; V = 39, p_uncorrected_ = 0.021, p_bonferroni_ = 0.042, β difference = −0.68, Cohen’s *d* = 0.4274], but not for hV5/MT+ [one-tailed Wilcoxon signed rank test; V = 92, p_uncorrected_ = 0.460, p_bonferroni_ = 0.920, β difference = −0.04, Cohen’s *d* = 0.0230](Figures 3B and C). In addition, applying TMS over left PMd and right hV5/MT+ did not extinguish the serial dependence effect, as slopes for previous trial speed were still lower than 0 [left PMd: one-tailed one sample Wilcoxon signed rank test: V = 24, p_uncorrected_ = 0.003, p_bonferroni_ = 0.012, β = −1.063, Cohen’s *d* = −0.771; right hV5/MT+: V = 4, p_uncorrected_ < 0.001, p_bonferroni_ < 0.001, β = −1.824, Cohen’s *d* = −1.333] (Figure 3B and C).

Applying TMS over Vertex also decreased serial dependence compared with the No-TMS condition [one-tailed Wilcoxon signed rank; V = 5, p < 0.001, β difference = −1.79, Cohen’s *d* = 0.8459](Figure 3B and C), but did not abolish the serial dependence effect [one-tailed one sample Wicoxon signed rank; V = 5, p_uncorrected_ < 0.001, p_bonferroni_ < 0.001,β = −1.864, Cohen’s *d* = −1.467]. The reduction was possibly due to confounder effects of TMS such as tactile sensation, noise and alertness (de Graaf and Sack 2011).

We next asked whether TMS also affected processing on the current trial, independent of the serial dependence (memory trace) examined above. In other words, did TMS affect temporal error and participants’ response variability independently of previous trial speed? To test this, we first performed a Friedman test on the current trial speed slopes of the multiple linear regression across conditions. Results indicated a difference in slope across conditions [χ (3) = 9.35, p = 0.025, Kendall’s W = 0.183], but comparing PMd and hV5/MT+ against Vertex did not show evidence for a difference [one-tailed Wilcoxon signed rank tests; left PMd: V = 46, p_uncorrected_ = 0.090, p_bonferroni_ = 0.180, β difference = −1.546, Cohen’s *d* = 0.469; right hV5/MT+: V =, p_uncorrected_ = 0.029, p_bonferroni_ = 0.058, β difference = −1.264, Cohen’s *d* = 0.497]. Second, we performed a Friedman test of temporal error across conditions, shown in Figure 4. The analysis did not show evidence for difference across conditions [χ (3) = 4.84, p = 0.184, Kendall’s W = 0.072]. However, we note that the temporal error for TMS over left PMd (33.78 ± 80.72, mean ± standard deviation) trended somewhat higher in comparison to the other three conditions (No-TMS: 17.40 ± 83.56; Vertex: 16.43 ± 85.23; right hV5/MT+: 18.34 ± 81.49, mean ± standard deviation). There was also no evidence for a difference in response variability (i.e. standard deviation of temporal error) across stimulation conditions [χ (3) = 5.40, p = 0.145; No-TMS: 54.74 ± 21.43; Vertex: 50.48 ± 25.64; right hV5/MT+: 51.16 ± 25.40; left PMd: 52.14 ± 24.30, mean ± standard deviation](Figure 4).

**Figure 4.**
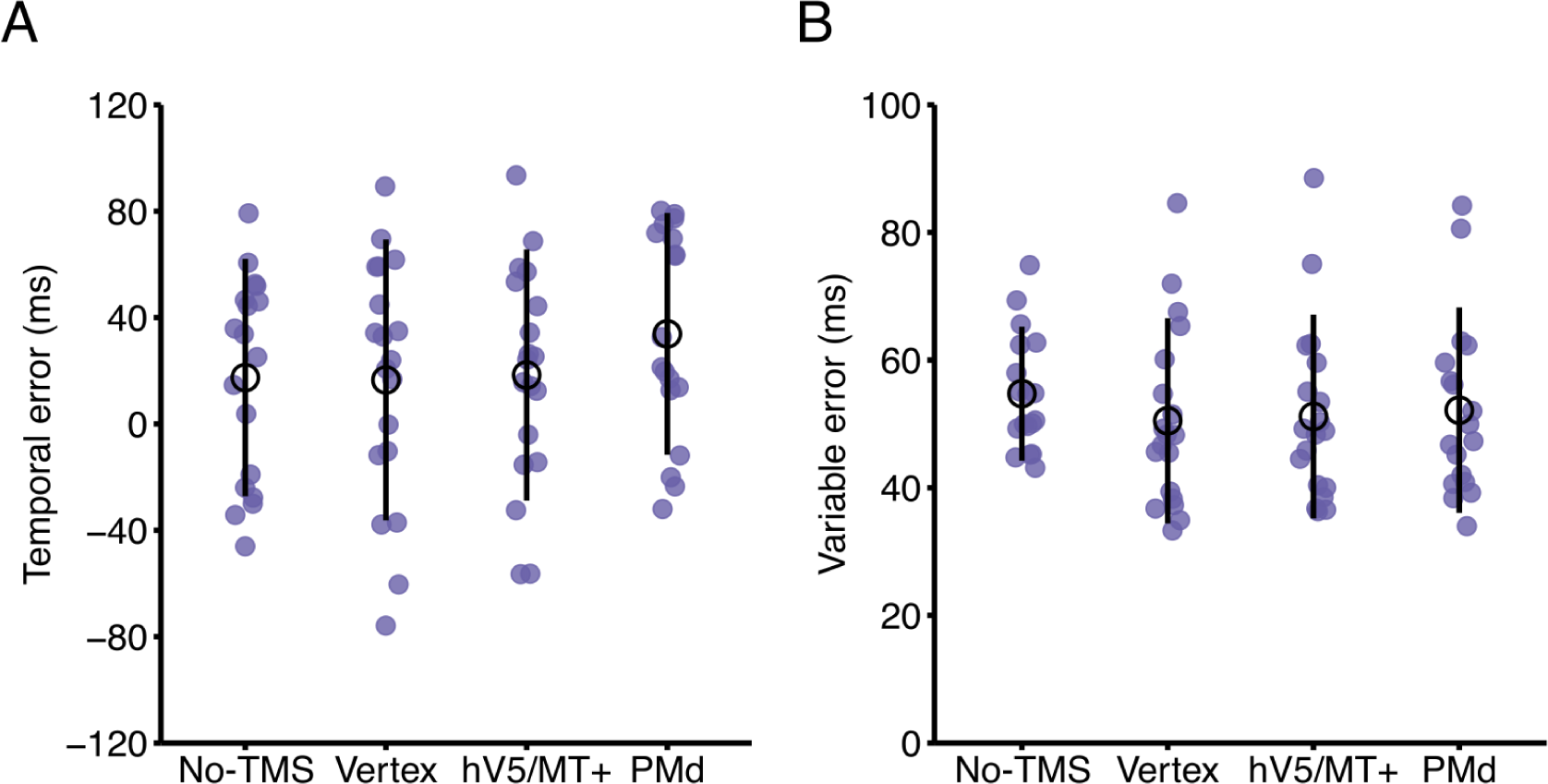
Absence of evidence for a TMS effect on the current trial apart from serial dependence. (A) Temporal error (ms) and (B) variable error (ms) averaged over previous and current trial speeds on No-TMS, Vertex, right hV5/MT+ and left PMd conditions. Blue circles represent the average temporal and variable errors for each participant. Black open circles represent the median temporal and variable errors within conditions. Error bars represent interquartile range within condition.

### Eye-tracker control results

Participants were able to maintain gaze around the fixation cross throughout the experiment and their gaze behavior did not differ across conditions (percentage of time participants gaze was kept 2° visual angle around fixation cross: F(3,39) = 0.3829, p = 0.7658, η^2^= 0.0286; number of saccades: F(3,39) = 0.0838, p = 0.9684, η^2^ = 0.0064).

## Discussion

Here we examined if serial dependence relies on an implicit short-term memory trace at cortical areas associated with visual motion processing or planning motor actions in a coincident timing task. We tested this prediction by applying TMS over right hV5/MT+ and left PMd in-between trials to disrupt the neural memory trace that putatively keeps information from one trial to the next (van de Ven and Sack 2013). We found that TMS over left PMd decreased serial dependence in comparison to stimulating Vertex. On the other hand, TMS over right hV5/MT+ had no effect compared to Vertex.

Apart from reducing serial dependence, we found no evidence that TMS had any other effect on performance, as the average temporal and variable errors did not differ across stimulation conditions. These results (i) corroborate the proposal that cortical regions encoding specific features of a task can store such information in short-term memory (c.f. Christophel et al. 2017), (ii) extend this proposal by showing the distributed storage of short-term memory occurs even when previous information is no longer needed to perform the task, and (iii) show that serial dependence causally relies on normal functioning of at least one such region, namely PMd.

### Location and nature of implicit short-term memory

For previous trial information to bias current behavior, it has to be stored and subsequently blended with incoming sensory information, motor plans or decisions. But what features of a task are stored in short-term memory and where is it stored? It has been proposed that the same regions that process and encode features of tasks or stimuli can also store those features in short-term memory (Christophel et al. 2017; Postle et al. 2015; Serences et al. 2016; Pasternak and Greenlee 2005; Kristijanson and Campana 2010). Our results corroborate this proposal, since TMS over one of these regions, left PMd, decreased serial dependence. Our results show that this mechanism is not restricted to perceptual explicit (e.g. Jerde et al. 2012; Riggall and Postle 2012) or perceptual implicit (e.g. Campana et al. 2002) tasks, but a general property of the neocortex that is also present in visuomotor integration tasks, and is invoked even in a task where participants are never asked to remember the features of the target (Fornaciai and Park 2018).

There is still debate of whether the short-term memory trace that causes serial dependence comes from sensory and perceptual (St. John-Saaltink et al. 2016; Makin et al. 2008; Cicchini et al. 2017; Fischer and Whitney, 2014; Manassi et al. 2018; Fornaciai and Park 2018) or post-perceptual processes (Bae and Luck 2020; Fritsche and de Lange, 2017; Kim et al. 2020; Pascucci et al. 2019). To avoid confusion, we are addressing the debate about whether the short-term memory trace is carried from sensory, perceptual, decision or motor processes *from* the previous trial, not whether the serial dependence effect on the current trial takes place during perceptual or post-perceptual stages of information processing (e.g. Fritsche and de Lange, 2017; Bliss et al. 2017; van Bergen and Jehee 2019; Bae and Luck 2019) since the current experimental design cannot address the latter. In the present study, applying TMS over PMd reduced serial dependence, whereas applying TMS over right hV5/MT+ had no effect (or one that did not differ from Vertex stimulation). These results suggest that the short-term memory trace influencing serial dependence relied on the neural substrate from the action performed by the individual in the previous trial.

Previous studies have shown the role of motor areas in explicit (Jerde et al. 2012) and implicit (Campana et al. 2007; O’Shea et al. 2007) short-term memory. In the present study, we show that PMd contralateral to movement planning and execution stores information implicitly. But what task or stimulus features are encoded in left PMd? The simplest explanation would be that the information that causes serial dependence comes from planning and executing the interception on the previous trial. This idea is in agreement with recent studies suggesting that the attractive bias is more likely to be caused by previous decisions and responses (Pascucci et al. 2019).

However, the bias could also be caused by temporal information stored from the previous movement or stimulus (e.g. Wiener et al. 2014). Alternatively, it is possible that PMd stores information about the speed of the previous trial, since premotor areas have been associated with perceiving visual information (Schubotz and von Cramon 2002a, 2002b; Schubotz 2007; Stadler et al. 2012). In addition, previous studies have shown that the specific motor action performed on the previous trial had a smaller influence in the current trial bias when compared to the perceptual decision (Cicchini et al. 2017), suggesting that the perturbed memory trace in our experiment might not be purely motor. Our results do not allow ruling in favor of any of these possibilities and future studies are needed to address these issues. In this context it should also be noted that growing evidence points to a direct oscillatory coupling between motor and visual sensory regions, potentially necessitating a more integrated viewpoint of the two systems (Benedetto et al. 2020; Benedetto et al., 2020).

In humans, the putative location of PMd is very close to that of the Frontal Eye Fields (FEF; c.f. Connolly, 2007), PMd usually being identified more dorsal than FEF within the superior frontal sulcus. The euclidean distance between FEF according to the neurosynth meta-analytic tool (Yarkoni et al. 2011) and the average left PMd across participants in our experiment is 3.4 mm (MNI; FEF: X = -28, Y = -4, Z = 52; PMd: X = -26, Y = -4.48, Z = 54.73). Given that the proximity between these regions is in the same order as the spatial precision of TMS stimulation, we might have stimulated FEF as well as PMd at the same time. FEF is a region implicated in controlling the gain of visuomotor transmission for smooth pursuit eye-movements (Tanaka and Lisberger, 2001). Although participants were able to keep their gaze at the fixation cross in both our localizer and main tasks, the oculomotor system receives a continuous drive of visual motion information from visual motion sensitive areas in the visual cortex (Lisberger 2015). Given that the smooth pursuit system might help predicting the speed and direction of moving objects necessary to perform interceptive actions (Bennett and Benguigui 2010; Foolken et al. 2021), it is possible that applying TMS over left premotor cortex disrupted the gain of visuomotor transmission stored in FEF in between trials, which reduced serial dependence. In that line, previous studies in monkeys have shown ramping activity in FEF neurons that are modulated by previous trial target speed: the greater the speed of the target on the previous trial, the stronger the ramping activity before the trial begins and the greater the initial eye-movement speed (Darlignton et al. 2018). In their modeling work, Darlington et al. (2018) suggest that such history effect is caused by storing previous trial information through an increase in synaptic weights of excitatory recurrent neurons. In our experiment, the further away the location of PMd was from meta-analytically derived FEF coordinates for each participant, the greater the serial dependence effect tended to be (Spearman rho = -0.45, p = 0.058), which is suggestive of a potential role also of FEF in our task. Whether FEF or PMd, we provide causal evidence that disrupting the memory trace in the premotor cortex decreases serial biases.

Stimulating right hV5/MT+ did not affect serial dependence compared to stimulating Vertex. Does it mean that hV5/MT+ does not keep information about previous trial information in an interceptive action? Although it might be the case and in agreement with recent findings suggesting that serial dependence is primarily biased by previous decisions and responses (Pascucci et al. 2019), we offer alternative explanations. A possible explanation is that the memory trace in hV5/MT+ is resistant to or protected against TMS stimulation. TMS effects depend heavily on the current state of the targeted neuronal population (Silvanto et al. 2008). In explicit short-term memory tasks, items held outside of the focus of attention present an activity pattern different from items held under the focus of attention (Lewis-Peacock and Postle 2012; LaRocque et al. 2013; Wolff et al. 2015, 2017; Rose et al. 2016). It could be the case that memory traces at the time we stimulated hV5/MT+ enter a state where stimulation does not affect it (Cattaneo et al. 2009; van de Ven et al. 2012; Pasternak and Zaskas, 2003). In support of this argument, applying TMS over hV5/MT+ in a working memory task about motion direction does not affect memory recall for items outside of the focus of attention, but does affect memory recall for items under the focus of attention (Zokaei et al. 2014). In addition, this apparent resistance to TMS stimulation may exist because both left and right visual regions might maintain a trace of the previous trial speed. Cortico-cortical connections between hV5/MT+ regions (Bridge et al. 2008; Genç et al. 2011), then, might make the memory trace more stable and resistant to TMS pulses. Lastly, other motion sensitive regions in the visual cortex (c.f. Orban et al. 2003) might have stored information about visual motion from one trial to the next. Not being able to address these issues is a limitation of the present study and the debate about whether the visual cortex can store information in short-term memory that leads to serial dependence remains open. Future investigations might address them by stimulating hV5/MT+ bilaterally, varying the strength of the stimulation pulse or its timing (Cattaneo et al. 2009; van de Ven et al. 2012; Pasternak and Zaskas, 2003), or targeting other visual motion sensitive regions. However, it is also noteworthy that numerous prior TMS studies targeting V5/MT+ yielded behavioral effects when TMS was applied unilaterally while visually stimulating bilaterally (Brascamp et al. 2008; Hotson and Anand 1999; Laycock et al. 2007; Silvanto et al. 2008), demonstrating that unilateral TMS can consistently affect motion perception of central motion displays.

### Serial dependence relies on short-term memory precision

How did TMS decrease the serial dependence effect? In virtual lesion experiments, high-intensity and high-frequency rTMS is usually associated with an increase in neuronal facilitation (Rossi et al. 2009), and we assumed this facilitation would disrupt the memory trace by introducing noise into the memorandum. In line with recent studies, we speculate that the network keeps the memory trace through short-term facilitation of synapses of neurons that have been activated to encode previous trial information (Mongilo et al. 2008; Kilpatrick 2018; Bliss and D’Esposito 2017). By applying TMS, we might be facilitating all synapses in the network at once, decreasing any specific tuning of such a memory trace. This decrease in network tuning for a specific stimulus, from a population coding perspective (see Pouget et al. 2013 for a review), could be interpreted as increasing the uncertainty of the memory trace. If the brain performs a weighted combination of previous and current information relying on memory and stimulus uncertainty (van Bergen and Jehee, 2019; Darlington et al. 2017; Darlington et al. 2018 Deravet et al. 2018; Kalm and Norris, 2018; Cicchini et al. 2018; Kwon and Knill 2013; Orban de Xivry et al. 2013), an increase in memory trace uncertainty would lead to a decrease in serial dependence (although see Barbosa et al. 2020 for an increase in serial dependence after applying TMS over prefrontal cortex in a working memory task). Future modeling and experimental work on short-term memory mechanisms and TMS effects on brain networks would be key to explaining such interactions.

Serial dependence is a ubiquitous phenomenon across different sensory modalities (for a review, see Kiyonaga et al. 2017). The mechanism that gives rise to it is still under investigation, but a few components seem to be fundamental for its emergence: the brain needs to keep previous trial information in short-term memory and blend it with incoming information (van Bergen and Jehee 2019; Kalm and Norris 2018; Chiccini et al. 2018; Kwon and Knill 2013). Here, we present evidence that premotor areas, more specifically PMd, have a potential role in storing previous trial information in implicit short-term memory in a visuomotor integration task, and that this information is responsible for causing biases on ongoing behavior. These results corroborate the perspective that areas associated with processing information of a stimulus or task also participate in maintaining that information in short-term memory (Postle 2015; Serences 2016; Christophel et al. 2017) even when this information is no longer relevant for current behavior.

## Acknowledgments

This work was funded by the Centre for Integrative Neuroscience (Tübingen, Germany), German Excellence Initiative grant number EXC307 (Germany), by the Max-Planck Society (Germany), and by Coordenação de Aperfeiçoamento de Pessoal de Nível Superior (CAPES, Brazil; scholarship awarded to RMAN, award number 99999.006725/2015-05).

